# Environment-wide and epigenome-wide association study of adiposity in “Children of 1997” birth cohort

**DOI:** 10.1101/2022.09.12.507623

**Authors:** Jie V Zhao, Bohan Fan, Jian Huang, BJ Cowling, SL Au Yeung, Andrea Baccarelli, GM Leung, C Mary Schooling

## Abstract

**Background:** Increasing childhood adiposity is a global issue requiring potentially local solutions to ensure it does not continue into adulthood. We systematically identified potentially modifiable targets of adiposity at the onset and end of puberty in Hong Kong the most economically developed major Chinese city.

**Methods:** We conducted an environment-wide association study (EWAS) and an epigenome-wide association study of adiposity to systematically assess associations with body mass index (BMI) and waist-hip ratio (WHR) in Hong Kong’s population-representative “Children of 1997” birth cohort. Univariable linear regression was used to select exposures related to adiposity at ~11.5 years (BMI n≤7,119, WHR n=5,691) and ~17.6 years (n = 3,618) at Bonferroni-corrected significance, and multivariable linear regression to adjust for potential confounders followed by replication (n=308) and CpG by CpG analysis (n=286) at ~23 years. Findings were compared with evidence from randomized controlled trials (RCTs) and Mendelian randomization (MR) studies.

**Results:** At ~11.5 and ~17.6 years the EWAS identified 14 and 37 exposures associated with BMI, as well as seven and 12 associated with WHR respectively. Most exposures had directionally consistent associations at ~23 years. Maternal second-hand smoking, maternal weight, and birth weight were consistently associated with adiposity. Diet (including dairy intake and artificially sweetened beverages), physical activity, snoring, binge eating, and earlier puberty were positively associated with BMI at ~17.6 years, while eating before sleep was inversely associated with BMI at ~17.6 years. Findings for birth weight, dairy intake, binge eating, and possibly earlier puberty are consistent with available evidence from RCTs or MR studies We found 21 CpGs related to BMI and 18 to WHR.

**Conclusions:** These novel insights into potentially modifiable factors associated with adiposity at the outset and the end of puberty could, if causal, inform future interventions to improve population health in Hong Kong and similar Chinese settings.

**Funding:** This study was supported by the Health and Medical Research Fund Research Fellowship, Food and Health Bureau, Hong Kong SAR Government (#04180097). The DNA extraction was supported by CFS-HKU1.

## Introduction

With improving living standards and socioeconomic development, non-communicable chronic diseases pose a heavy burden on society in both developed and developing countries (1). Obesity is a well-established risk factor for multiple chronic diseases, including cardiovascular disease, diabetes, and cancer (2). Obesity has increased substantially in many settings, including Hong Kong. Obesity is complicated with multifactorial risk factors, such as socioeconomic position (SEP), mood disturbance and genetic factors (3), so there is a need to consider all factors holistically, and to evaluate their roles in adiposity comprehensively and systematically (4). Moreover, given most studies on targets of obesity are conducted in a western setting (3), a comprehensive assessment of modifiable factors of early life adiposity in a non-western setting, with a different social structure, provides a valuable opportunity to identify novel exposures.

Environment-wide association studies (EWAS) enable us to assess a variety of exposures across the human environmental exposome in a high-throughput manner (5), similar to genome wide association studies for genetic associations. Previous EWAS have been successfully performed on outcomes, such as type 2 diabetes (6), cardiovascular disease (7), and childhood obesity in western settings (8, 9) but not in a Chinese setting. In the previous EWAS of childhood adiposity in the US (6-17 years old), UK and Europe (6-11 years old), second-hand smoking was related to higher childhood BMI, whilst some other exposures, such as vitamins, were not consistently associated with adiposity in these settings (8, 9). Observational studies are open to confounding by SEP, thus assessing associations in a different social context can triangulate the evidence concerning early life adiposity.

In addition to the environmental factors, it is increasingly realized that epigenetic factors, which are also modifiable, may play an important role in adiposity (10). DNA methylation, which refers to the addition of a methyl group to the 5’ position of a cytosine residue of the DNA, is the most frequently examined epigenetic modification (11). DNA methylation may modulate gene expression and thereby influence susceptibility to obesity or obesity-related chronic disease (11). Epigenome-wide association study provides an approach to identify the related epigenetic loci in a comprehensive way. For example, DNA methylation at cg06500161 was previously identified as related to adiposity in the US (10). Unlike genetic variants, DNA methylation is modifiable and may change in response to environmental factors or disease and therefore might be open to confounding by these factors (11, 12). As such, findings from western settings may not be generalizable to Chinese populations.

In this situation, Hong Kong, a non-Western developed setting, can provide unique insights into health determinants. Most Chinese people in Hong Kong are first-, second- or third-generation migrants from the neighbouring province of Guangdong in southern China. Dietary habits of people in Hong Kong are similar to those in southern China, although also influenced by western culture (13). Lifestyle in Hong Kong also differs from more commonly studied western populations on some important attributes, for example, active smoking among Chinese mothers was rare while maternal exposure to second-hand smoking during pregnancy was common before the smoking ban in public and workplaces was implemented in 2007 (14). Moreover, most current theories concerning aetiology of and disparities in chronic diseases originate from observations in long-term developed populations of European descent. However, in Hong Kong the economic transition from pre- to post-industrial living conditions has occurred within one lifetime of the older people (15), whereas children today in Hong Kong represent the first generation of Chinese to grow up in a post-industrial Chinese setting; this is unrivalled anywhere in the world (16). As such, a study in young Chinese people in Hong Kong, a different setting provides “a sentinel for populations currently experiencing very rapid economic development” (17), may help identify whether these associations reflect SEP within a specific context or are biologically based as well as having the potential to identify any attributes relevant to the majority of the global population but not necessarily evident in more commonly studied Western populations, such as maternal birthplace (18). For example, in the “Children of 1997” birth cohort, a large cohort in Hong Kong, the associations of sugar-sweetened beverages (19), breastfeeding (20), milk consumption frequency (21), sleep duration (22) and parental smoking (23) with childhood and adolescent adiposity have been examined, with some important differences detected. The associations for breastfeeding in Hong Kong are much more similar to those seen in randomized controlled trials (RCTs) than those typically seen in western settings (20). To take advantage of this unique setting, we conducted an environment-wide and epigenome-wide association study, to identify further potential drivers of adiposity. We considered adiposity at the outset and at the end of puberty, because puberty involves a re-orientation from childhood priorities to adulthood (24), and exposures associated with adiposity at puberty may be important for health in later life (25, 26).

## Methods

### Subjects

The study takes advantage of the “Children of 1997” birth cohort, a large (n= 8,327) population-representative Chinese cohort in Hong Kong (16). The participants were originally recruited shortly after birth in April and May 1997 at all of the 49 Maternal and Child Health Centers (MCHCs) in Hong Kong, which provide free check-ups and immunizations. The study included 88% of births in the relevant period. A self-administered questionnaire in Chinese was used at baseline to collect information on family, education, birth characteristics, infant feeding, and second-hand smoke exposure. The initial study was designed to provide a short-term assessment of the effects of second-hand smoking and included follow-up via the MCHCs until 18 months. In 2005, funded by the Health and Health Services Research Fund (HHSRF) and Health and Medical Research Fund (HMRF) we extended the information on this cohort via record linkage to include infant characteristics, serious morbidity, childhood adiposity, pubertal development, history of migration and socio-economic position; with regular updates on subsequent growth obtained from the Student Health Service, including annual height and weight measurements from age ~6 years, and in this study, we used height and weight at age ~11.5 years. In 2007, with support from The University of Hong Kong University Research Committee Strategic Research Theme of Public Health, we instituted a program to re-establish and maintain direct contact with the cohort through direct mailing (newsletters, birthday cards and seasonal cards) and the mass media (press conference and a full-length television documentary). We have since conducted three questionnaires/telephone surveys and an in-person Biobank clinical follow-up at age ~17.6 years (Phase 1 in 2013-6 included 3,460 people with mean age 17.5 years and Phase 2 in the second half of 2017 included 158 people at mean age 19.5 years). Over 3,600 participants attended the Biobank clinical follow-up, with their blood samples stored (Appendix Figure 1). In 2020 (at age ~23 years), we conducted a follow-up survey to obtain updated information on anthropometric measurements. The flow chart of this study was shown in Appendix Figure 2.

### Assessment of adiposity

At ~11.5 years, BMI was calculated from height and weight measurements records provided by Student Health Service, the Department of Health. WHR was calculated based on waist and hip circumstance collected in Survey I conducted in 2008-2009. At ~17.6 years, in both phases of the Biobank clinical follow-up, BMI was assessed by bio-electrical Impedance Analysis (BIA) with a Tanita segmental body composition monitor (Tanita BC-545, Tanita Co., Tokyo, Japan). Waist and hip circumference measurements were made using a tape twice following a standard protocol by trained technicians and nurses. In the follow-up survey at ~23 years, questionnaires were sent to 700 participants randomly selected from those with blood samples available and with BMI below the 25^th^ centile or above the 75^th^ centile. The questionnaires were accompanied by clear instructions on anthropometric measurement and a tape measure (the same as used in the Biobank clinical follow-up). In total, 308 participants replied and provided their waist, hip, height, and body weight.

### Assessment of DNA methylation

DNA methylation was conducted in 288 participants randomly selected from the 308 participants in the follow-up survey. DNA were extracted from buffy coat samples previously stored at −80 degrees using EZI DNA blood kit (QIAGEN) with magnetic particle technology. DNA methylation was assessed using the Illumina Methylation EPIC Beadchip, which interrogated the methylation status of over 850,000 CpG sites. We conducted quality control using the “ewastools” package (27), which included an evaluation of control metrics monitoring the various experimental steps, such as bisulfite conversion or staining and a sex check comparing actual sex to the records. After sample-level quality control, we excluded two samples that had a sex mismatch, so 286 samples (168 women and 118 men) were included in the analysis. We corrected for dye bias using RELIC (28), without normalization. At the probe-level, we excluded non-CpG probes and probes located on the sex chromosomes; a total of 843,393 probes remained for analyses.

### Exposures and categorization

Appendix Table 1 shows exposure categorization and data sources. After excluding exposures with missing values ≥ 50%, we included 123 exposures for BMI and 115 exposures for WHR at ~11.5 years, and 441 exposures for BMI and WHR at ~17.6 years using information from the original study, record linkage, the three surveys, i.e., Survey I (2008-2009), Survey II (2010-2012) and Survey III (2011-2012), as well as the Biobank clinical follow-up (Appendix Figure 1). The exposures considered for adiposity at ~11.5 years were classified into 12 categories, including baseline characteristics, SEP, family history, paternal information, maternal information, infant feeding and caring, diet (measured at Survey I at ~11.5 years old), children’s health, parents’ health, physical activity, lifestyle, and home facilities and pets. The exposures at 17.6 years were classified into 16 categories: baseline characteristics, SEP, family history, paternal information, maternal information, infant feeding and caring, diet (measured at the Biobank clinical follow-up at ~17.6 years), children’s use of medications, children’s health, parent’s health, physical activity, home facilities and pets, moods and feelings, academic performance, sleep, and pubertal timing.

### Statistical analysis

In the EWAS, similar to genome-wide association studies (29), first we used univariable linear regression to assess associations of each of the exposures with the measures of adiposity at ages ~11.5 years and ~17.6 years. We conducted the analysis in people with both exposure and outcome available, specifically, in up to 7,119 participants for BMI at ~11.5 years, 5,691 participants for WHR at ~11.5 years, and 3,618 participants for BMI and WHR at ~17.6 years; their baseline characteristics are shown in Table 1. We only considered exposures reaching Bonferroni-corrected significance (for example, p < 0.05/441 = 1.2*10^-4^ for adiposity at ~17.6 years) to account for multiple testing (30). Second, we used multivariable linear regression controlling for potential confounders (sex, housing type at birth, household income at birth, maternal second-hand smoking during pregnancy, maternal age at birth, maternal education, maternal birth place, and the interaction of maternal education with maternal birthplace (18)) at age ~11.5 years and ~17.6 years, and excluded exposures that had over 50% of change-in-estimates ratios (31). Third, we replicated the associations for the selected exposures in the follow-up survey (n=308) at age ~23 years and compared the direction of associations with those at ~11.5 years and ~17.6 years. Associations with consistent directions of associations at ~11.5 years or ~17.6 years with those at ~23 years suggest a consistent association by age. Finally, exposures that remained after controlling for confounders, were compared with the evidence from existing RCTs and Mendelian randomization (MR) studies, a study design which uses genetic variants as instrument and provides less confounded associations (32).

**Table 1.**
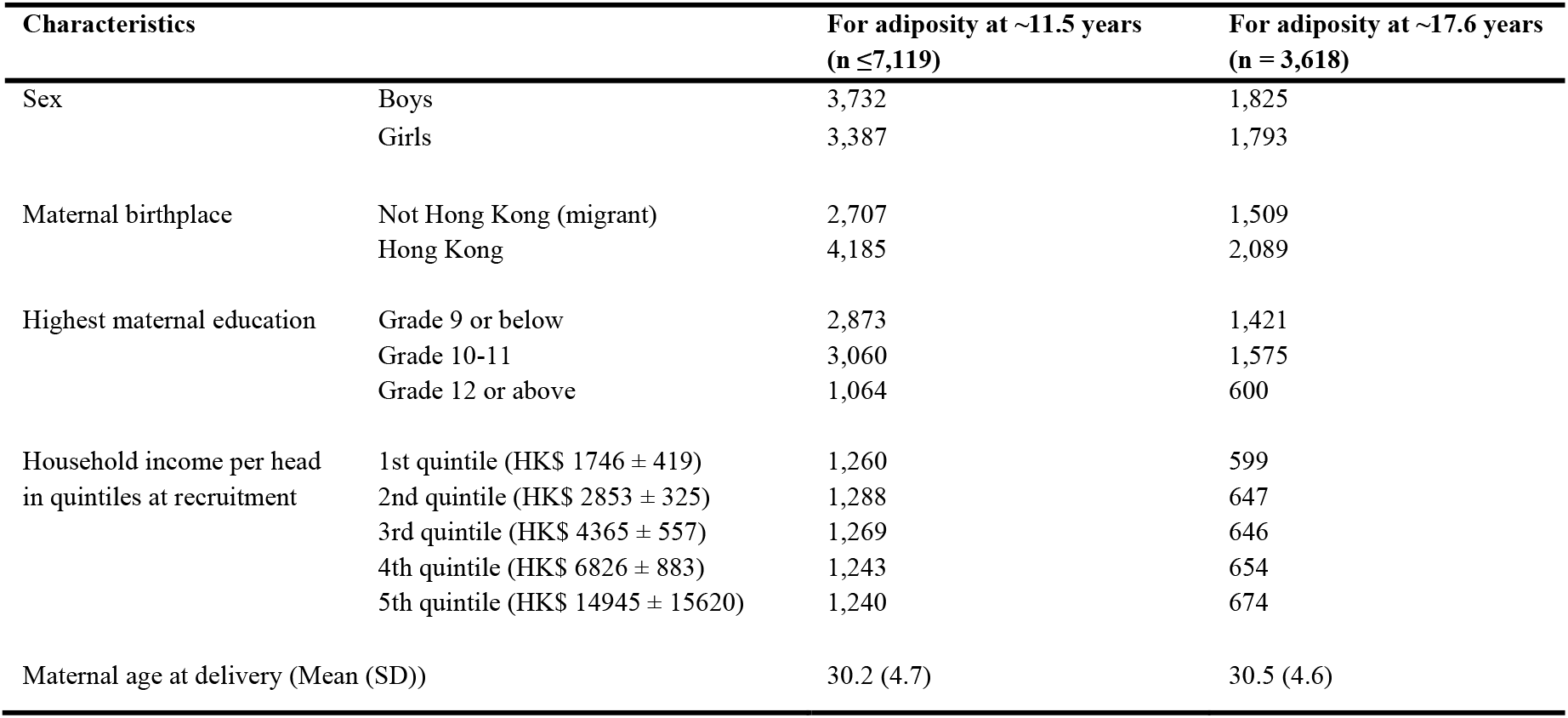
Baseline characteristics of Hong Kong’s “Children of 1997” birth cohort included in the EWAS of adiposity at ~11.5 years and ~17.6 years.

| Characteristics |  | For adiposity at ~11.5 years<br>(n ≤ 7,119) | For adiposity at ~17.6 years<br>(n = 3,618) |
| --- | --- | --- | --- |
| Sex | Boys | 3,732 | 1,825 |
|  | Girls | 3,387 | 1,793 |
| Maternal birthplace | Not Hong Kong (migrant) | 2,707 | 1,509 |
|  | Hong Kong | 4,185 | 2,089 |
| Highest maternal education | Grade 9 or below | 2,873 | 1,421 |
|  | Grade 10-11 | 3,060 | 1,575 |
|  | Grade 12 or above | 1,064 | 600 |
| Household income per head<br>in quintiles at recruitment | 1st quintile (HK\$ 1746 ± 419) | 1,260 | 599 |
| | 2nd quintile (HK\$ 2853 ± 325) | 1,288 | 647 |
| | 3rd quintile (HK\$ 4365 ± 557) | 1,269 | 646 |
| | 4th quintile (HK\$ 6826 ± 883) | 1,243 | 654 |
| | 5th quintile (HK\$ 14945 ± 15620) | 1,240 | 674 |
| Maternal age at delivery (Mean (SD)) |  | 30.2 (4.7) | 30.5 (4.6) |

In the epigenome-wide association study, considering the heteroscedasticity in methylation beta-values (33), we used robust linear regression models to assess the epigenome-wide association of each CpG with BMI and WHR at age ~23 years. We adjusted for age at blood draw for DNA methylation, age at follow-up survey, sex, cell type proportion, methylation assay batches, maternal second-hand smoking during pregnancy, maternal education, maternal birthplace, and household income at birth. The significance was considered as false discovery rate (FDR)<0.05 using a Benjamini Hochberg correction, given the relatively small sample size in the epigenome-wide association study. To estimate genomic inflation, we used a Bayesian method that estimates inflation more accurately in epigenome-wide association studies based on the empirical null distributions (34), implemented using the R package “bacon”.

## Results

Figures 1 and 2 showed the association of each exposure with BMI and WHR at ~11.5 years and ~17.6 years. At ~11.5 years, 18 associations with BMI and 19 associations with WHR remained after Bonferroni correction (Figure 1). Of these 18 associations with BMI, 14 associations with BMI remained after controlling for confounders (Table 2). Eleven exposures had concordant direction of associations with BMI at ~23 years (Table 3). Of the 19 associations with WHR at ~11.5 years, seven exposures remained after controlling for confounders (Table 4) and six had the same direction of association with WHR at ~23 years (Table 5). At ~17.6 years, 37 associations with BMI and 19 with WHR remained after Bonferroni correction (Figure 2). Of these 37 associations with BMI, all remained after controlling for confounders (Table 6), and 32 exposures had the same direction of association with BMI at ~23 years (Table 7). Of the 19 associations with WHR at ~17.6 years, 12 remained after controlling for confounders (Table 8), and ten had the same directions of association with WHR at ~23 years (Table 9).

**Figure 1.**
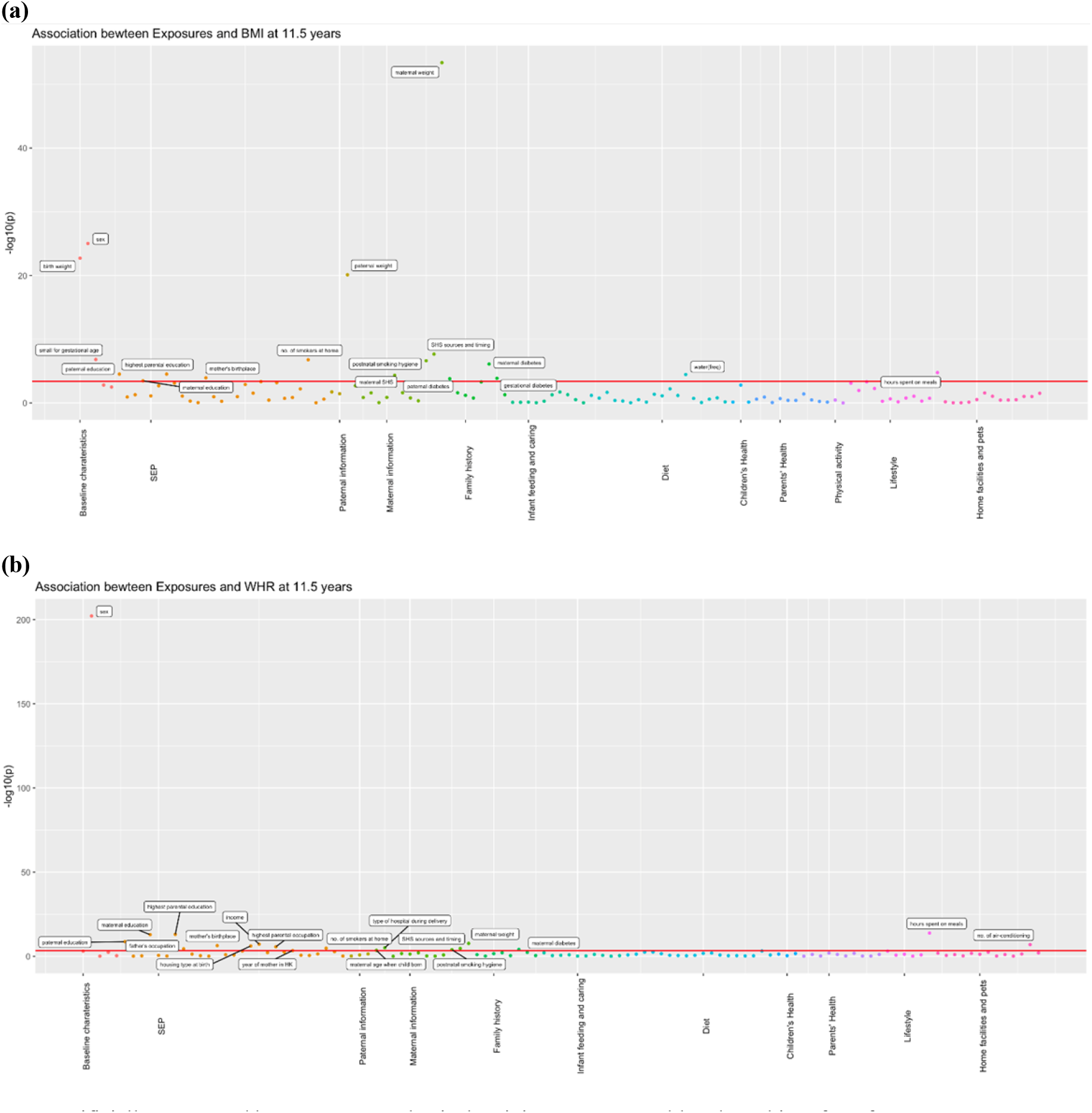
Associations of all exposures with BMI and WHR at age ~11.5 years in the univariable regression in Hong Kong’s “Children of 1997” birth cohort. ASB, artificially sweetened beverages; PA, physical activity; SHS, second-hand smoking; freq, frequency. In total, we included 123 exposures for BMI at 11.5 years and 115 exposures for WHR at 11.5 years. The cut-off lines indicate Bonferroni corrected p thresholds (p < 0.05/123 = 4.07 *10^-4^ for BMI, p < 0.05/115 = 4.35*10^-4^ for WHR).

**Figure 2.**
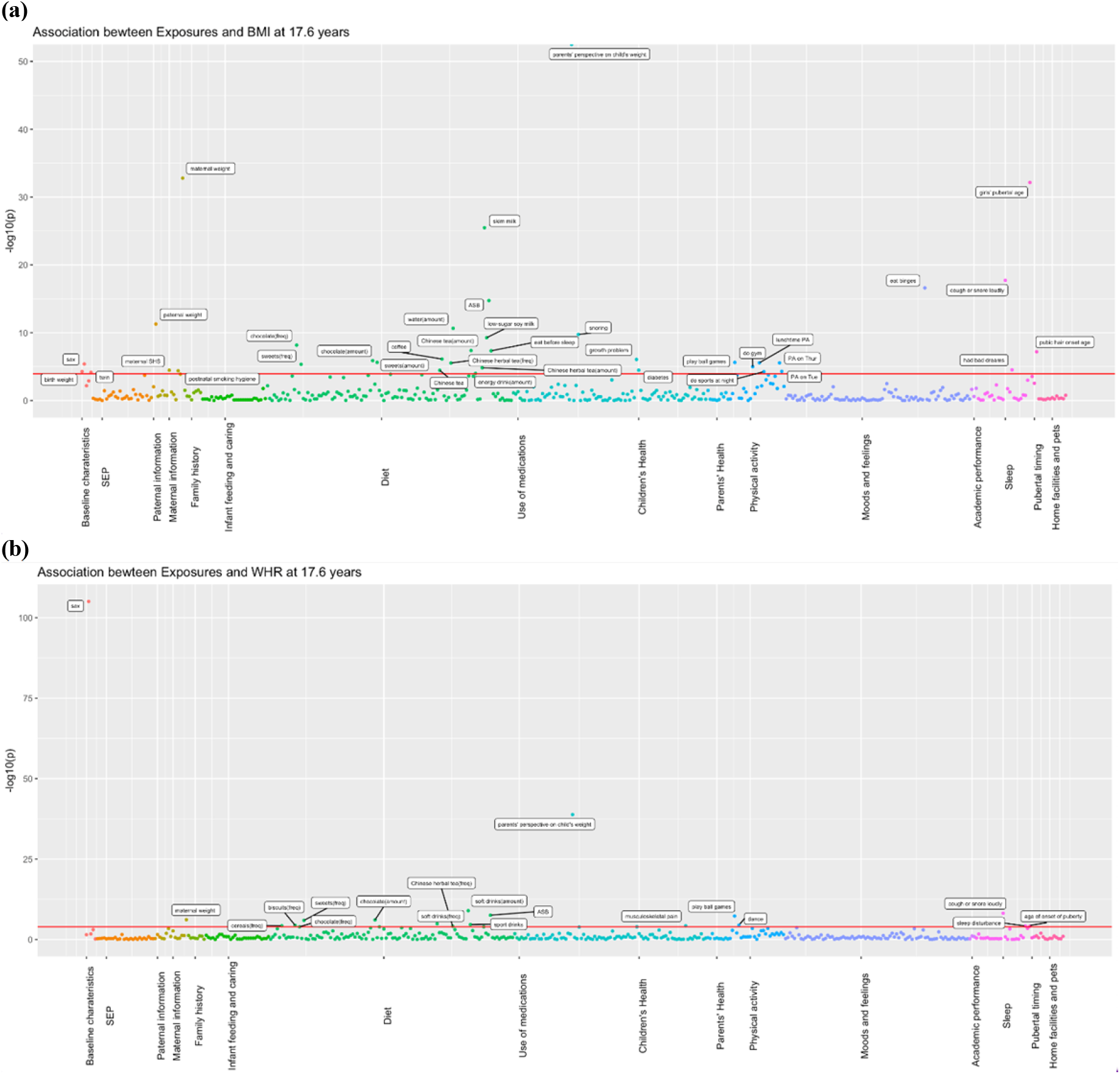
Associations of all exposures with BMI and WHR at age ~17.6 years in the univariable regression in 3,618 participants of Hong Kong’s “Children of 1997” birth cohort in the Biobank clinical follow-up. ASB, artificially sweetened beverages; PA, physical activity; SHS, second-hand smoking; freq, frequency The cut-off lines indicate Bonferroni corrected p thresholds (p < 0.05/441 exposures= 1.2*10^-4^ for BMI and WHR).

**Table 2.** Associations of selected exposures with BMI at age ~11.5 years in up to 7,119 participants of Hong Kong’s “Children of 1997” birth cohort after controlling for confounders.

| Group Names | Variable description | Beta | p value |
| --- | --- | --- | --- |
| Baseline characteristics | birth weight | 0.40 | 5.40E-15 |
| Baseline characteristics | sex | 0.85 | 5.65E-22 |
| Baseline characteristics | small for gestational age: birth weight<10% by sex and gestational week distribution in singletons | -0.68 | 9.93E-06 |
| SEP | no. of smokers at home | 0.15 | 2.83E-02 |
| Paternal information | paternal weight | 0.05 | 1.58E-18 |
| Maternal information | maternal weight | 0.09 | 1.84E-49 |
| Maternal information | postnatal smoking hygiene | 0.10 | 7.79E-05 |
| Maternal information | second-hand smoke by sources and timing | 0.08 | 2.69E-05 |
| Maternal information | mother exposed to second-hand smoke during pregnancy | 0.15 | 4.07E-03 |
| Family history | paternal diabetes | 0.69 | 5.82E-03 |
| Family history | maternal diabetes | 1.45 | 2.47E-05 |
| Family history | gestational diabetes | 0.68 | 2.28E-04 |
| Diet | water: frequency of consumption in the last week | 0.32 | 9.45E-06 |
| Lifestyle | having meals: hours spent yesterday | -0.40 | 2.06E-03 |

postnatal smoking hygiene refers to second-hand smoking by timing (pre- and/or postnatal)

**Table 3.**
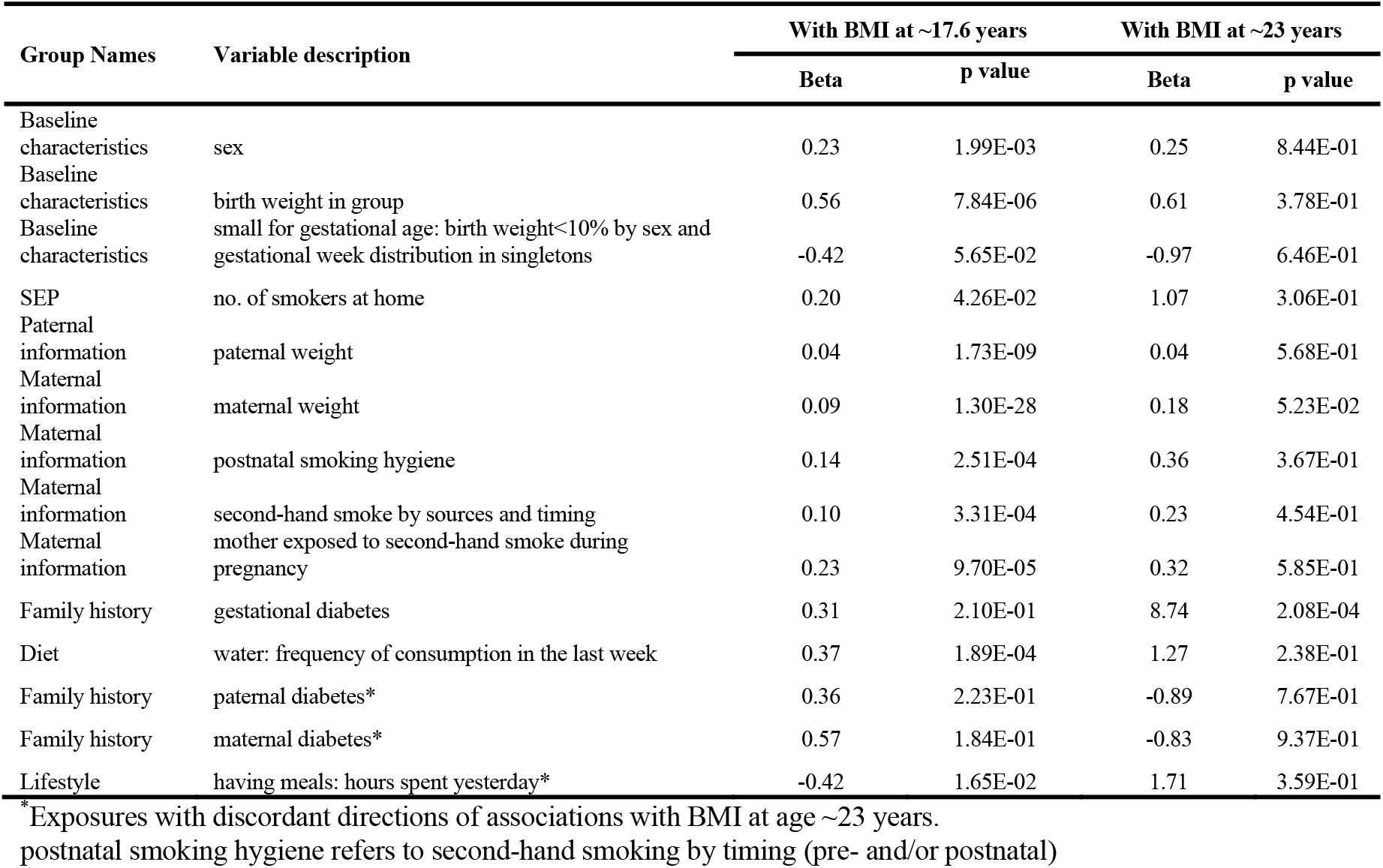
Associations of selected exposures for BMI at age ~11.5 years with BMI at age ~17.6 years (n=3,618) and ~23 years (n=308) in participants from Hong Kong’s “Children of 1997” birth cohort after controlling for confounders.

**Table 4.** Associations of selected exposures with WHR at age ~11.5 years after controlling for confounders in 5,691 participants of Hong Kong’s “Children of 1997” birth cohort.

| Group Names | Variable description | Beta | p value |
| --- | --- | --- | --- |
| Baseline characteristics | sex | 0.05 | 6.58E-171 |
| SEP | housing type at birth (home ownership) | -0.01 | 2.70E-02 |
| SEP | maternal education level | -0.01 | 4.90E-05 |
| SEP | mother's birthplace | -0.01 | 1.18E-01 |
| Maternal information | maternal weight | 0.00 | 7.98E-10 |
| Family history | maternal diabetes | 0.03 | 2.30E-05 |
| Lifestyle | having meals: hours spent yesterday | -0.01 | 8.15E-09 |

**Table 5.** Associations of selected exposures for WHR at age ~11.5 years with WHR at age ~17.6 years (n=3,618) and ~23 years (n=308) in participants from Hong Kong’s “Children of 1997” birth cohort after controlling for confounders.

| Group Names | Variable description | With WHR at ~17.6 years |  | With WHR at ~23 years |  |
| --- | --- | --- | --- | --- | --- |
|  |  | Beta | p value | Beta | p value |
| Baseline characteristics | sex | 0.04 | 1.28E-88 | 0.082 | 4.30E-18 |
| Maternal information | maternal weight | 0.001 | 1.13E-06 | 0.001 | 6.58E-02 |
| SEP | housing type at birth (home ownership) | -0.001 | 8.51E-01 | -0.01 | 4.20E-01 |
| SEP | maternal education level | -0.004 | 2.03E-01 | -0.001 | 9.02E-01 |
| SEP | mother's birthplace | -0.01 | 2.76E-01 | -0.006 | 7.97E-01 |
| Lifestyle | having meals: hours spent yesterday | -4.73E-03 | 1.30E-01 | -0.002 | 8.11E-01 |
| Family history | maternal diabetes* | 0.01 | 1.27E-01 | -0.017 | 7.82E-01 |

\*Exposures with discordant directions of associations with WHR at age ~23 years.

**Table 6.**
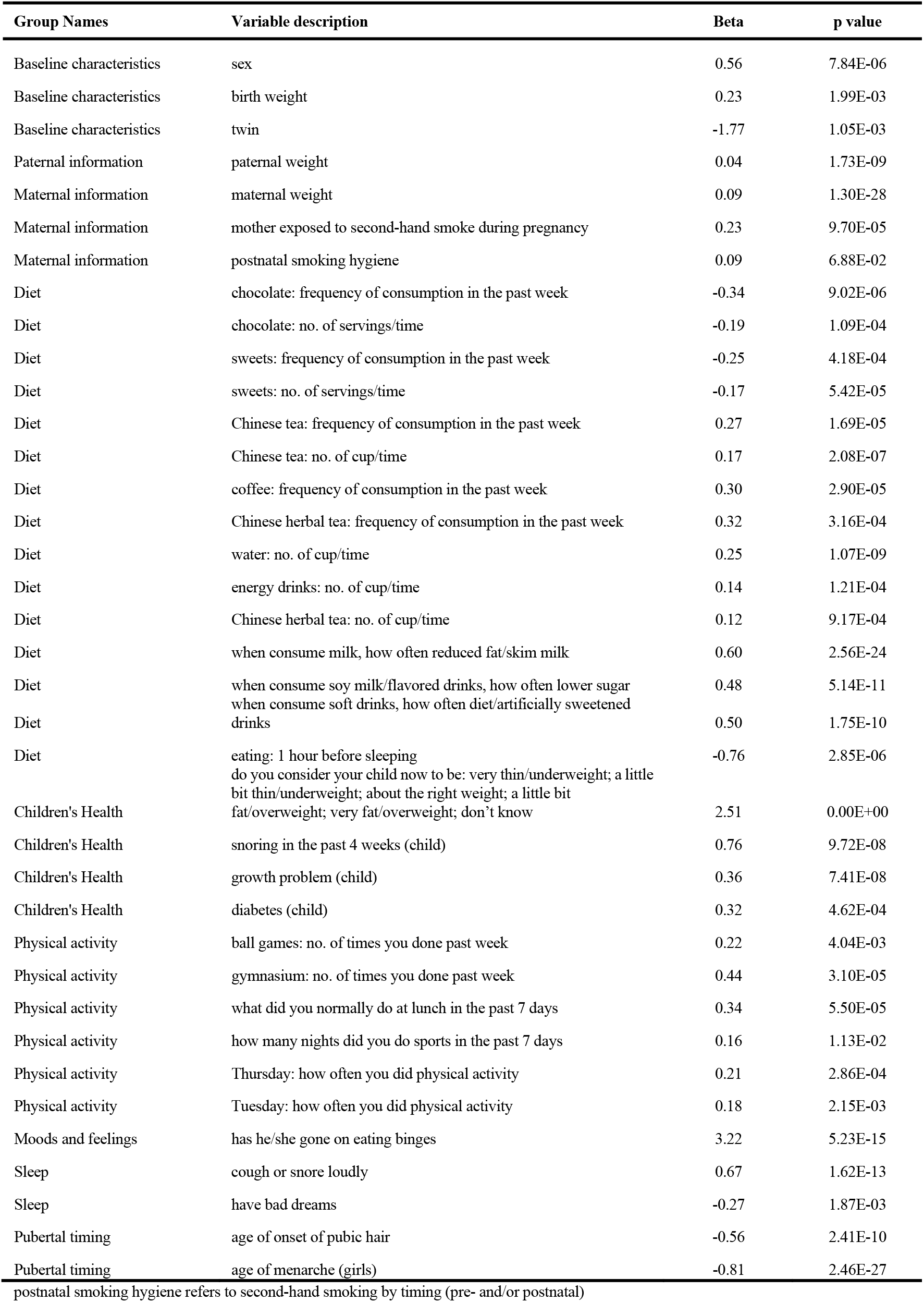
Associations of selected exposures with BMI at age ~17.6 years after controlling for confounders in 3,618 participants of Hong Kong’s “Children of 1997” birth cohort in the Biobank clinical follow-up.

**Table 7.**
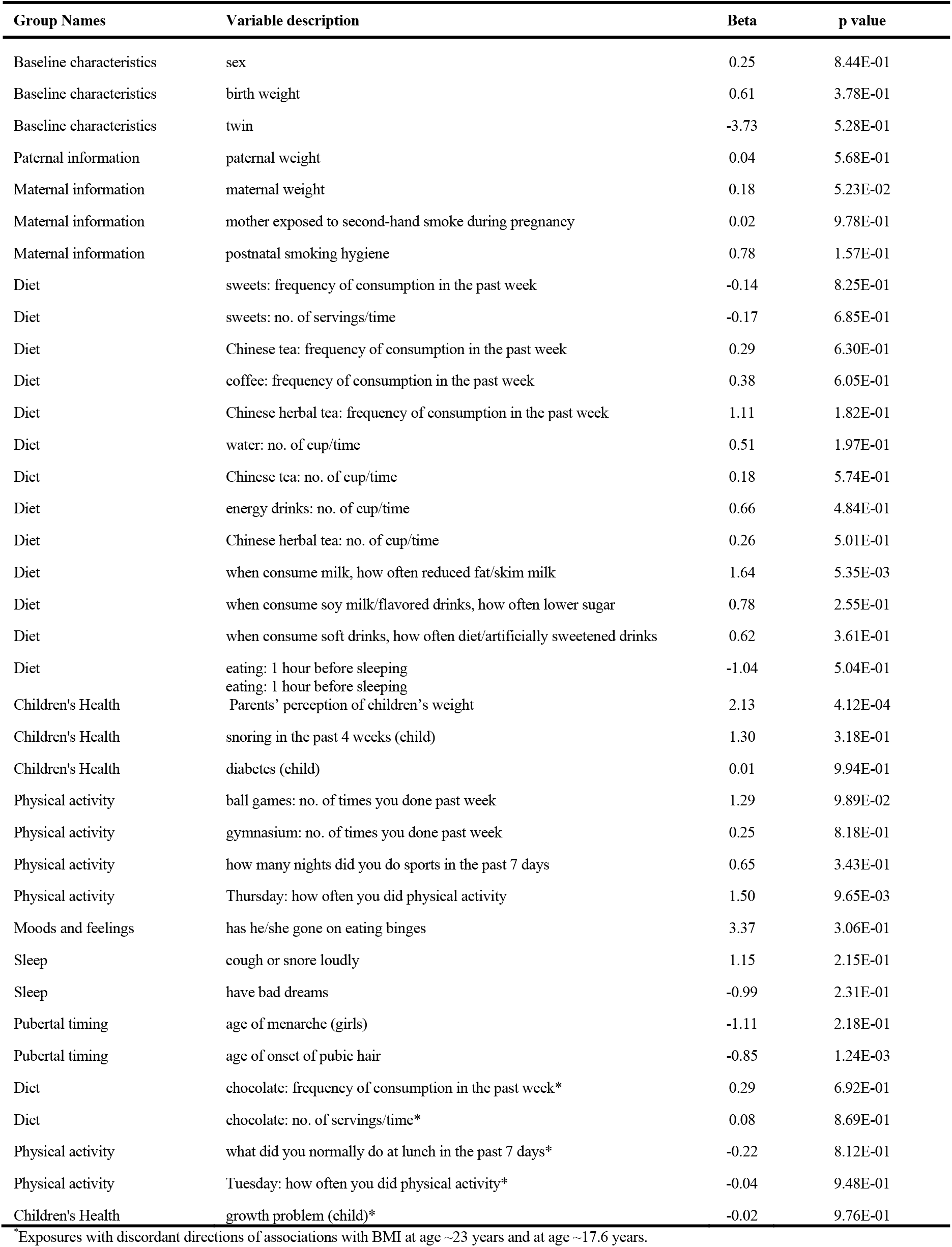
Associations of selected exposures for BMI at age ~17.6 years with BMI at age ~23 years in participants from Hong Kong’s “Children of 1997” birth cohort in the follow-up survey after controlling for confounders (n=308)

**Table 8.**
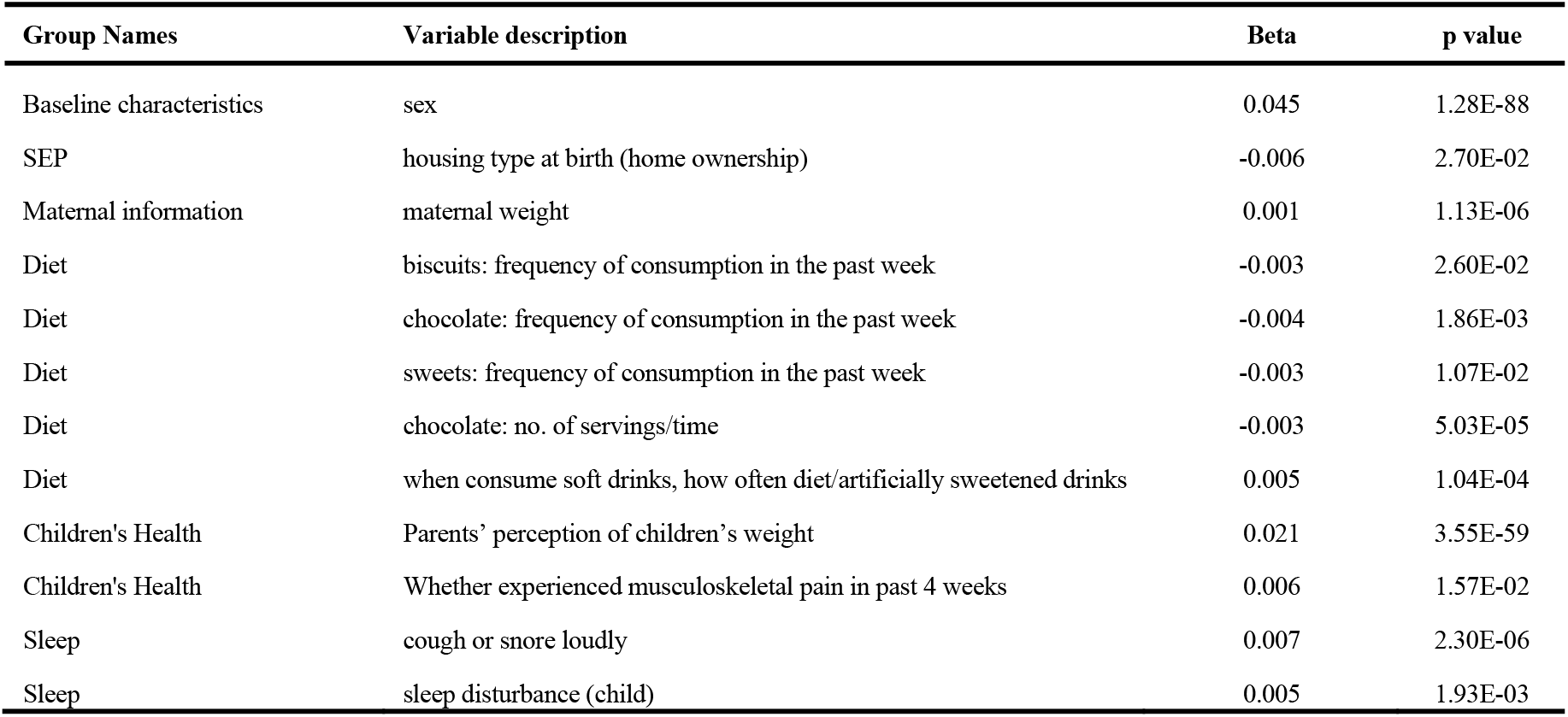
Associations of selected exposures with WHR at age ~17.6 years after controlling for confounders in 3,618 participants of Hong Kong’s “Children of 1997” birth cohort in the Biobank clinical follow-up.

| Group Names | Variable description | Beta | p value |
| --- | --- | --- | --- |
| Baseline characteristics | sex | 0.045 | 1.28E-88 |
| SEP | housing type at birth (home ownership) | -0.006 | 2.70E-02 |
| Maternal information | maternal weight | 0.001 | 1.13E-06 |
| Diet | biscuits: frequency of consumption in the past week | -0.003 | 2.60E-02 |
| Diet | chocolate: frequency of consumption in the past week | -0.004 | 1.86E-03 |
| Diet | sweets: frequency of consumption in the past week | -0.003 | 1.07E-02 |
| Diet | chocolate: no. of servings/time | -0.003 | 5.03E-05 |
| Diet | when consume soft drinks, how often diet/artificially sweetened drinks | 0.005 | 1.04E-04 |
| Children's Health | Parents' perception of children's weight | 0.021 | 3.55E-59 |
| Children's Health | Whether experienced musculoskeletal pain in past 4 weeks | 0.006 | 1.57E-02 |
| Sleep | cough or snore loudly | 0.007 | 2.30E-06 |
| Sleep | sleep disturbance (child) | 0.005 | 1.93E-03 |

**Table 9.**
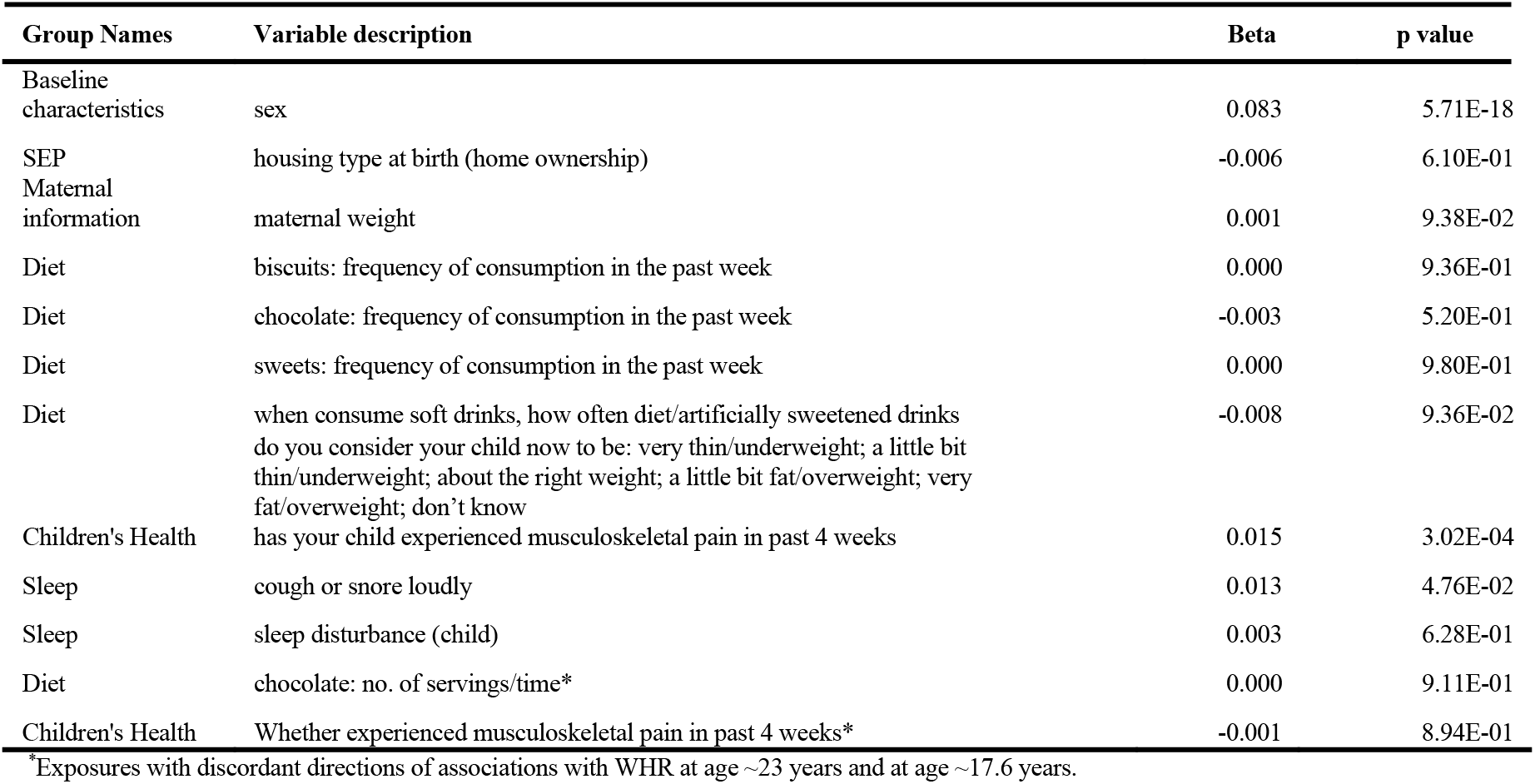
Associations of selected exposures for WHR at age ~17.6 years with WHR at age ~23 years in participants from Hong Kong’s “Children of 1997” birth cohort in the follow-up survey after controlling for confounders (n=308)

Specifically, for adiposity at ~11.5 years, we found sex (being male), higher birth weight, maternal second-hand smoking, higher parental weight, family history of diabetes, gestation diabetes, and more water consumption were associated with higher BMI, while being small for gestational age and spending more time having meals were associated with lower BMI at ~11.5 years (Table 2). However, the associations for family history of diabetes and time spent on meals showed inconsistent directions of associations for BMI ~23 years (Table 3). Regarding WHR, the associations were generally consistent with those seen for BMI (Table 4) and showed consistent directions of associations with those at ~17.6 years and ~23 years, except for maternal diabetes (Table 5).

For adiposity at ~17.6 years, in addition to some shared factors, including sex, birth weight, parental weight, and maternal second-hand smoking, we found some aspects of diet (i.e., more artificially sweetened beverage (ASB), lower-sugar soy milk, reduced-fat/skim milk, Chinese herbal tea, Chinese tea, energy drinks, coffee, and fish consumptions), physical activity, health (i.e., diabetes, growth problem and snoring), earlier puberty and binge eating associated with higher BMI at ~17.6 years (Table 6). Being a twin, sweets and chocolate consumption, eating before sleep, and having bad dreams were associated with lower BMI at ~17.6 years (Table 6). Most exposures had the same direction of association at ~23 years (Table 7). However, the association of chocolate consumption with BMI at ~23 years was in the other direction (Table 8). Regarding WHR, the associations were generally consistent with those seen for BMI. Sex, drinking ASB, children’s health, and coughing or snoring during sleep were also related to WHR at ~17.6 years (Table 9) and ~23 years (Table 9).

Regarding the comparison with RCTs and MR studies, we found RCTs on dark chocolate consumption (35), water consumption promotion (36), and physical activity (37), and MR studies related to drinking coffee (38), dairy intake (39), binge eating (40), physical activity (41), snoring (42), puberty (43, 44), birth weight (45), and maternal adiposity (46, 47) (Table 10). The available evidence from both RCTs and MR studies show that more physical activity lowers BMI (37), and MR studies show that dairy intake (39), binge eating (40), and earlier puberty (43, 44) are associated with higher BMI.

**Table 10.** Evidence from published systematic reviews, RCTs and MR studies regarding the role of exposures selected in our EWAS in obesity.

| Exposure | Published RCTs studies | Published MR studies |
| --- | --- | --- |
| Water consumption | Intervention on promoting water consumption showed no effect on BMI.<sup>38</sup> | NA |
| Reduced-fat/ skim milk consumption | NA | MR study on different types of milk consumption (i.e., reduced-fat, skimmed, reduced-sugar milk) among children is lacking. Nevertheless, one MR study suggested higher dairy intake was associated with higher adult's BMI.<sup>41</sup> |
| Coffee consumption | NA | Genetically predicted more coffee intake was not associated with obesity, BMI, or waist circumference in two large adult population cohorts.<sup>40</sup> Also, most MR studies do not support a role of caffeine consumption on BMI or waist circumference.<sup>40,61</sup> |
| Chocolate consumption | Meta-analysis of RCTs did not support a significant effect of cocoa/dark chocolate supplementation on body weight or BMI.<sup>37</sup> | NA |
| Diabetes | NA | MR study supported genetic predisposition to higher childhood BMI was associated with risk of type 2 diabetes.<sup>78</sup> |
| Binge eating | NA | The MR study suggests a bi-directional association, i.e., more binge eating and overeating are associated with higher BMI in later life, and higher children's BMI is associated with more binge eating.<sup>42</sup> |
| Physical exercises | A systematic review shows school based RCTs targeting physical activity, or physical activity combined with diet interventions, were effective in reducing BMI among children.<sup>39</sup> | Genetically predicted more physical activity was associated with lower BMI.<sup>43</sup> |
| Snoring | NA | MR study suggests genetically predicted BMI is related to snoring.<sup>44</sup> |
| Pubertal timing | NA | Genetically predicted earlier age at puberty was related to higher BMI,<sup>45,46</sup> but pleiotropy might exist.<sup>45</sup> It is also possible that genetically predicted BMI was associated with earlier age at puberty.<sup>69</sup> |
| Birth weight | NA | MR analyses indicated positive causal associations of birthweight with BMI in the UKB.<sup>47</sup> |
| Maternal adiposity during pregnancy | NA | Using BMI polygenic risk scores calculated from maternal non-transmitted alleles, previous MR studies using mother-offspring pairs from two large UK cohorts did not support causal associations between maternal pre/early pregnancy BMI and offspring adolescent adiposity.<sup>48,49</sup> |

In the epigenome-wide association study of 286 participants we identified 21 CpGs for BMI at ~23 years in the genes *RBM16, SCN2B, AGPAT4, TFCP2, SLC24A4, TECPR2, KSR1, RPTOR, GTF3C3, ZNF827, TXNDC15, C2*, and *RPS6KA2* and 18 for WHR in the genes *LANCL2, C6orf195, MIR4535, CTRL, LYRM9, DCDC2, DIRC3, RPS6KA2, LPP, NFIC, MIR7641-2, ZNF141, RNF213, OPA3*, and *RRS1* (Tables 11-12 and Figures 3-4). cg14630200 in *RPS6KA2* was a shared CpG for both BMI and WHR. The genetic inflation factors (lambda) were 1.006 and 1.005 respectively.

**Table 11.** Associations of CpGs with BMI at age ~23 years in 286 participants from Hong Kong’s “Children of 1997” birth cohort in the follow-up survey.

| CpG | gene | Beta | p value |
| --- | --- | --- | --- |
| cg00379266 | *RBM16* | -69.7 | 4.6E-02 |
| cg01525498 | *RPTOR* | -40.2 | 1.3E-04 |
| cg02200725 | - | -18.9 | 3.7E-02 |
| cg04563671 | *SCN2B* | 14.8 | 1.7E-02 |
| cg07243141 | - | 24.6 | 4.6E-02 |
| cg08205320 | - | -21.9 | 4.0E-03 |
| cg09639369 | *SLC24A4* | -15.7 | 3.7E-02 |
| cg09655876 | *AGPAT4* | -17.3 | 4.6E-02 |
| cg11297723 | *TFCP2* | -38.9 | 4.6E-02 |
| cg11342941 | *AGPAT4* | -11.6 | 4.6E-02 |
| cg11937362 | - | 109 | 4.6E-02 |
| cg12484266 | *C2* | 83 | 1.0E-03 |
| cg14630200 | *RPS6KA2* | -45.1 | 3.0E-06 |
| cg14850190 | *KSR1* | 103.2 | 3.7E-02 |
| cg15145978 | - | -20.4 | 4.6E-02 |
| cg18681028 | - | -48.5 | 6.4E-10 |
| cg18893311 | - | 30.8 | 1.0E-02 |
| cg22197830 | *TXNDC15* | -68.8 | 3.7E-02 |
| cg25356423 | *ZNF827* | -14 | 1.7E-02 |
| cg25903177 | - | -31.4 | 3.7E-02 |
| cg26347007 | *GTF3C3* | -45.8 | 4.6E-02 |

CpGs reaching significance of FDR<0.05 were shown in the table.

**Table 12.** Associations of CpGs with WHR at age ~23 years in 286 participants from Hong Kong’s “Children of 1997” birth cohort in the follow-up survey.

| CpG | gene | Beta | p value |
| --- | --- | --- | --- |
| cg00905170 | *RRS1* | 2.04 | 4.8E-02 |
| cg00952960 | *LANCL2* | -0.54 | 2.5E-05 |
| cg04405211 | *C6orf195* | 0.35 | 4.8E-02 |
| cg05059349 | - | 0.26 | 2.6E-02 |
| cg07416310 | *DCDC2* | -0.45 | 9.0E-03 |
| cg09422806 | *NFIC* | 0.5 | 5.0E-03 |
| cg09684856 | *DIRC3* | -0.79 | 1.3E-02 |
| cg09907395 | *RNF213* | 1.09 | 1.7E-02 |
| cg11344771 | *LPP* | -0.79 | 1.8E-03 |
| cg14630200 | *RPS6KA2* | -0.46 | 3.3E-02 |
| cg17007012 | *MIR4535* | -0.35 | 2.8E-05 |
| cg17043713 | *ZNF141* | 0.55 | 9.0E-03 |
| cg17769811 | *CTRL* | -0.76 | 4.0E-04 |
| cg19481727 | *MIR7641-2* | -0.36 | 3.3E-02 |
| cg19579176 | - | -0.35 | 4.8E-02 |
| cg22364817 | - | -1.46 | 3.7E-02 |
| cg25097095 | *LYRM9* | -0.41 | 2.0E-03 |
| cg27384074 | *OPA3* | 0.55 | 1.4E-02 |

CpGs reaching significance of FDR<0.05 were shown in the table

**Figure 3.**
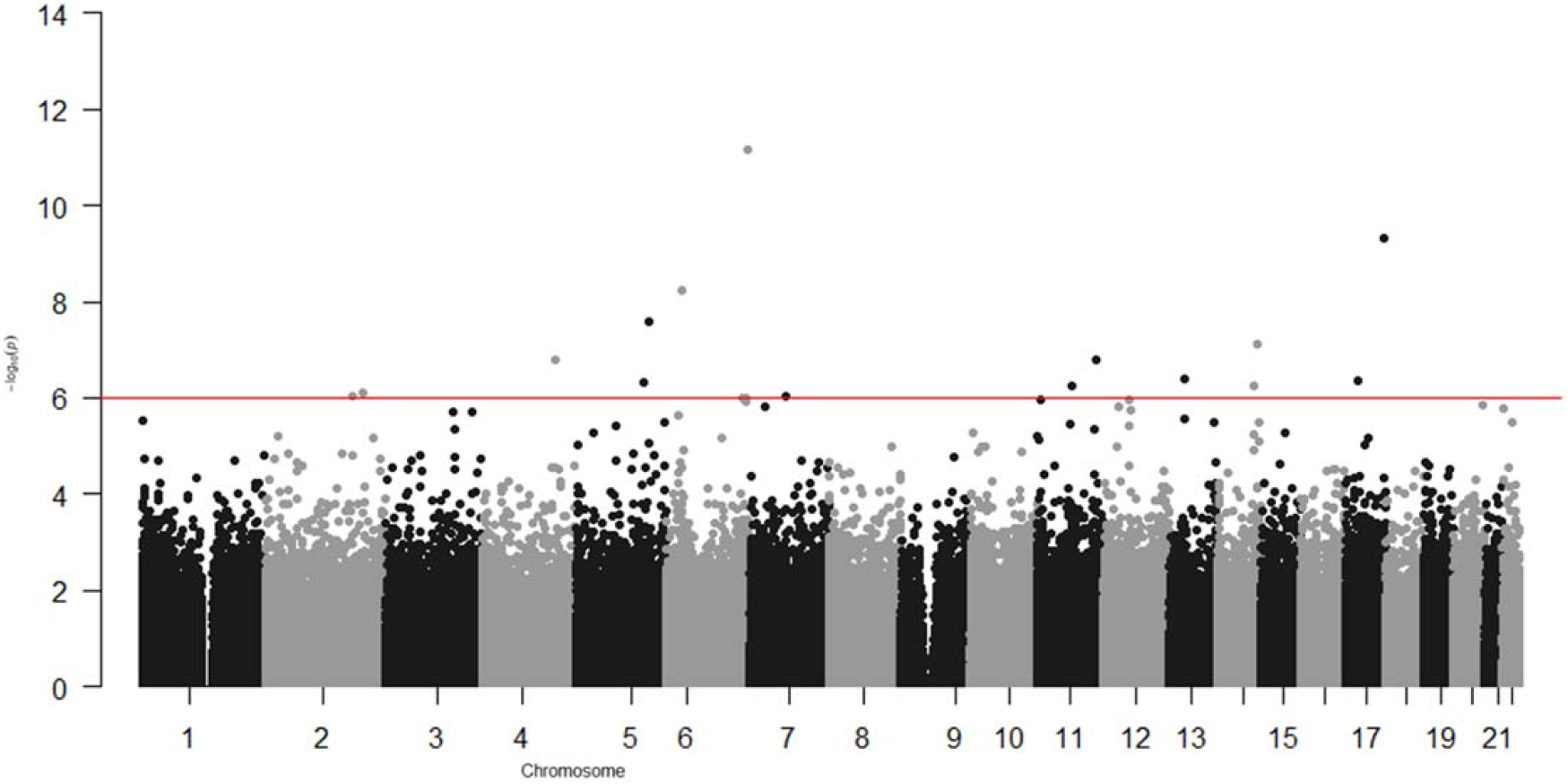
Epigenome-wide association with BMI at age ~23 years in 286 participants from Hong Kong’s “Children of 1997” birth cohort in the follow-up survey.

**Figure 4.**
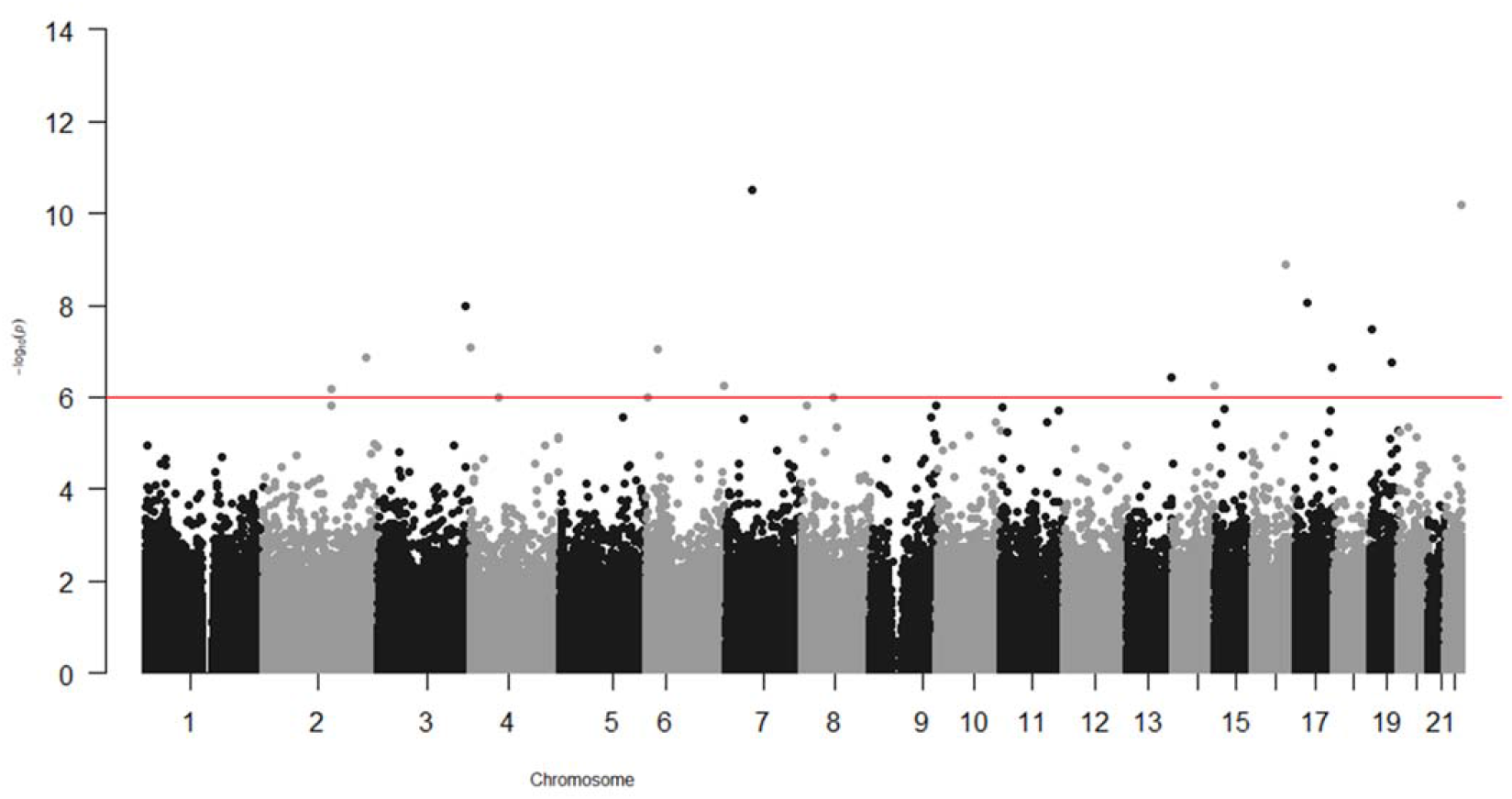
Epigenome-wide association with WHR at age ~23 years in 286 participants from Hong Kong’s “Children of 1997” birth cohort in the follow-up survey.

## Discussion

In this environment-wide and epigenome-wide association study, we systematically examined associations of over four hundred exposures with adiposity in a unique Chinese birth cohort, as well as the association of DNA methylation with adiposity. Building on the previous studies in this birth cohort (19–21, 23), we not only confirmed established risk factors, such as maternal second-hand smoking (48), but also added by identifying novel exposures not reported in previous EWAS in western settings (8, 9), such as consumption of ASB and soymilk. The comparison with RCTs or MR studies support a role of higher birth weight, dairy intake, binge eating and possibly earlier puberty in adiposity. We also identified several CpGs related to BMI and WHR in young Chinese, as reported in other populations (49).

Our study found that maternal second-hand smoking was consistently associated with adiposity at different ages, which is consistent with the concerns repeatedly raised in previous studies (23, 48), and adds support to the policy of banning smoking cigarettes and alternative smoking products in all indoor areas including workplaces and public places, as well as certain outdoor areas, such as open areas of schools, leisure facilities, bathing beaches, and public transport facilities in Hong Kong (50). Maternal weight is another maternal factor related to higher BMI and WHR consistently at different ages before adulthood. However, recent MR studies do not support a role of maternal overweight in offspring obesity (46, 47). Gestational diabetes was also identified to be associated with adiposity at ~11.5 years, and the positive association remained for adiposity at ~17.6 years and ~23 years. It would be worthwhile to test its role in MR studies.

Regarding dietary factors, as the dietary assessments were more comprehensively conducted in Biobank clinical follow up, the identified dietary factors were mainly for adiposity at ~17.6 years. Interestingly, we found that children who consume more ASB have higher BMI, which is consistent with meta-analyses of cohort studies (51, 52). Consistent with a previous study in this birth cohort (19), we did not find an association of sugar-sweetened beverages with adiposity. The different associations for ASB and sugar-sweetened beverages might be because few consumed sugar-sweetened beverages regularly (6.8% consumed daily) (19), while many consumed ASB (43% participants reported consumption). Consistent with our EWAS, an EWAS in the US also found that consumption of aspartame, a synthetic non-nutritive sweetener, was positively associated with abdominal obesity (53). ASB intake may induce appetite for similar sweet foods, leading to excess energy intake (54). The consistency across settings suggests this association is less likely to be confounded. However, whether it can be used as a target of intervention needs to be tested in a randomized, placebo-controlled trial (52).

Another interesting finding is that milk consumption was not related to adiposity at ~11.5 years, while reduced-fat/skim milk consumption was associated with higher BMI at ~17.6 years, with a consistent direction of associations for BMI at ~23 years. Our findings are consistent with a previous study in this cohort, which showed milk consumption frequency was not associated with BMI at 13 years (21), however, the previous study did not assess the specific type of milk. Our findings are different from a cross-sectional study in Portugal which shows more skimmed or semi-skimmed milk consumption was associated with lower abdominal obesity (55). An explanation is that the observed associations of reduced-fat or skim milk with higher BMI could be due to increased muscle mass rather than body fat mass, or residual confounding by SEP. Alternatively, it might be due to reverse causality as young people with higher BMI might be more motivated to consume a specific diet. Interestingly, our findings are more consistent with an MR study suggesting genetically predicted higher dairy intake was associated with higher BMI (39).

We also found tea or coffee consumption were associated with higher BMI at ~17.6 years. RCTs of coffee or tea consumption are scarce among children and adolescents because they require long-term adherence. MR studies do not suggest that coffee consumption affects adiposity (38, 56), so the observed association might be due to confounding or chance. Similarly, the associations of chocolate and sweets intake with lower BMI at ~17.6 years, as well as physical activity with higher BMI at ~17.6 years are not consistent with RCTs of dark chocolate consumption (35) and physical activity (37)), or MR studies of physical activity (41), and might be due to confounding or reverse causality.

Echoing the increasing attention to the role of mood and emotion in obesity control (3), we found that binge eating was associated with higher BMI at ~17.6 years, with a consistent direction of association in the follow-up, consistent with an MR study (40). Our findings are also in line with the National Institute for Health and Care Excellence (NICE) guidance which also included binge eating in the consideration of children’s weight management (57). The underlying mechanism has not been clarified, but in general mental wellbeing may be linked to adiposity via the neurohormonal weight control network concerning the hypothalamus (58, 59) as well as via psychosocial factors, lifestyle and behaviour.

As regards lifestyle, consistent with previous observational studies (22), we found sleep might play a role in childhood adiposity at ~17.6 years. Despite a lack of RCTs, MR findings suggest sleep deprivation may be a causal factor for obesity (22, 60). Our study also shows coughing or snoring at night had a positive association with childhood BMI and WHR, which has not been identified in earlier EWAS (8, 9). MR studies suggest genetically predicted BMI is positively associated with snoring (61), while the association of snoring with BMI is less clear. As such, we cannot exclude the possibility of reverse causality in our observation.

Consistent with previous studies (62), we found that earlier pubertal age for girls was related to higher subsequent BMI at ~17.6 years. Consistently, a previous study in this birth cohort suggested that maternal age at puberty was associated with offspring BMI at puberty (63). This finding is also consistent with MR studies showing genetically predicted earlier age at puberty related to higher BMI (43, 44) although we cannot exclude a relation in the other direction (64). Clarifying the bi-directional association would be worthwhile in future studies.

In the epigenome-wide association study, we found DNA methylation at *RPS6KA2* was associated with both BMI and WHR, consistent with the previous epigenome-wide association studies of obesity in different populations (49). Our study also identified several other genes, such as *ZNF827, MIR7641-2, RAPTOR, KSR1, GTF3C3* and *NFIC*, whose role in obesity or obesity-related disorders has been consistently shown in previous studies. For example, *ZNF827, MIR7641-2* and *RAPTOR* have been reported to be related to obesity (65, 66) and/or overweight (67). *KSR1* has been reported to be related to the regulation of glucose homeostasis (68), and *GTF3C3* related to obesity-related dysglycaemia (69). *NFIC*, which encodes nuclear factor I-C, regulates adipocyte differentiation (70)*. Opa3*, a novel regulator of mitochondrial function, controls thermogenesis and abdominal fat mass (71). The consistency of our study with other studies in different settings with different confounding structures suggests these association are less likely to be a product of confounding.

### Strengths and Limitations

To our knowledge, this study is the first study comprehensively assessing environmental factors related to adiposity at the outset and at end of puberty in Asians, including some exposures specifically relevant in Asians, such as soymilk intake. We also replicated the associations in a follow-up survey and compared our findings with those from studies of different designs. Nevertheless, several limitations exist. First, the sample size for the EWAS and epigenome-wide association study is relatively small. Replication in a larger study is needed. Second, misclassification is possible for the exposures, which typically biases towards the null (72). The use of questionnaires to ascertain exposures is prone to recall bias and social desirability bias, however, some previous studies within this cohort have suggested accurate reporting (23). To minimize these possible biases, exposures measurements were collected using standard protocols and equipment with clear instructions. Finally, despite the minimal level of confounding in this cohort for some key exposures, such as breastfeeding, residual confounding likely exists, so the associations are not definitive.

### Implications

In this study, we not only confirmed established risk factors, such as maternal second-hand smoking, but also identified several factors not reported or not examined in previous EWAS in western countries (8, 9), such as ASB consumption and soymilk intake. The comparison with RCTs or MR studies support a role of dairy intake, binge eating and possibly earlier puberty, suggesting these factors or their drivers (for example, sex hormones as the drivers of age at puberty) might be considered as potential targets for intervention. Other factors, such as soymilk intake, need to be tested in RCTs. We also identified several methylation loci related to adiposity. Our study, in comparison with previous studies in different settings and studies with different study designs, provides potential drivers of adiposity, applicable to Hong Kong Chinese, with relevance to health policy interventions and future research.

## Conclusions

This study takes advantage of the unique setting of Hong Kong and provides more insight about the role of environmental exposures and epigenetics in early life adiposity. If these associations are found to be causal, they may provide novel intervention targets to improve population health.

## Ethical approval

Ethical approval for this study, including the follow-up survey at ~23 years and comprehensive health-related analyses, was obtained from the University of Hong Kong-Hospital Authority Hong Kong West Cluster, Joint Institutional Review Board, Hong Kong Special Administrative Region, China (Reference numbers: UW13-367; UW19-367).

## Declaration

None declared.

## Acknowledgements

This study was supported by the Health and Medical Research Fund Research Fellowship, Food and Health Bureau, Hong Kong SAR Government (#04180097). The DNA extraction was supported by CFS-HKU1. We would like to sincerely thank Food and Health Bureau for funding this project. We would also like thank all the participants and research staff in this project.

## Data availability

The data used in this study are based on the “Children of 1997” Birth cohort, maintained by the School of Public Health, The University of Hong Kong. With the approved ethics for this study, the individual participant data cannot be made freely available online. Interested parties can access the data used in this study upon reasonable request, with approval by the birth cohort team. As part of this process, researchers will be required to submit a project concept for approval, to ensure the data is being used responsibly, ethically, and for scientifically sound projects. Requesters should be employees of a recognized academic institution, health service organization, or charitable research organization with experience in medical research. Requestors should be able to demonstrate, through their peer-reviewed publications in the area of interest, their ability to carry out the proposed study. Source data files have been uploaded for each of the results figures (Figures 1 to 4) showing the model summary data for plotting the Manhattan plots in environment-wide and epigenome-wide associations with BMI and WHR. Source code for the analyses has been uploaded as Source Code.

## Reporting

The study conforms with the STROBE checklist, which was attached as a supplementary file.

