## Appendix Figure 1 for "Environment-wide and epigenome-wide association study of adiposity in “Children of 1997” birth cohort"

### Appendix Figure 1. Timeline of “Children of 1997” Birth Cohort

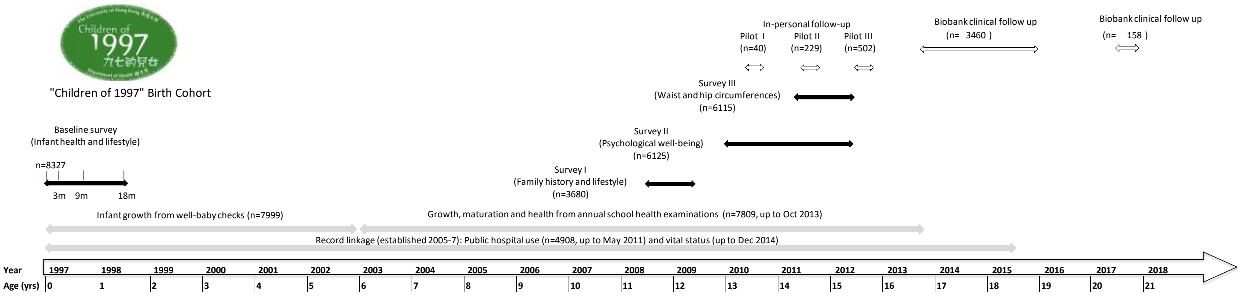


*Response rate for 3 months, 9 months and 18 months follow-up: 82%, 83% and 74%
*Number of respondents to all 3 surveys (Survey I,II,III): 3246(39%)
*Number of respondents to any of the 3 surveys: 6628 (80%)
*In-person follow-up includes anthropometric measurements, blood pressure, blood test, lung function test, biomaterials (such as stool, urine, hair and toenail samples) and etc.

Pilot study I(Feb 2010-Apr 2010): n=40
Pilot study II(Jun 2011-Aug 2011):n=229
Pilot study III (Jun 2012-Aug 2012):n=502
Biobank (Aug 2013-Jan 2016):n=3460

*Number of permanent withdrawal as of January 2018: 29
